# Nanoscale fracture of defective popgraphene monolayer

**DOI:** 10.1101/463505

**Authors:** Fanchao Meng, Ming Ni, Feng Chen, Jun Song, Dong Wei

## Abstract

A new carbon allotrope, namely popgraphene, has been recently demonstrated to possess high potentials for nanodevice applications. Here, the fracture of defective popgraphene was studied using molecular dynamics simulations and continuum modeling. Three scenarios of defects were considered, including an individual point defect, distributed point defects, and nanocracks. It was found that the fracture stress of popgraphene with an individual point defect was governed by both the geometry of the defect and the critical bond where fracture initiates. Moreover, the fracture stress of popgraphene with distributed point defects was discovered to be inversely proportional to the defect density, showing a nice linear trend. Furthermore, for popgraphene with a nanocrack, it failed in a brittle fashion and exhibited a negligible lattice trapping effect. Griffith criterion was subsequently employed with the consideration of crack deflection to accurately predict the dependence of fracture stress on crack size. The present study lays a mechanistic foundation for nanoscale applications of popgraphene and offers a better understanding of the roles of defects in fracture of low-dimensional materials.

## 1 INTRODUCTION

The remarkable properties^1- 3^ and promising applications^4- 6^ offered by two-dimensional (2D) graphene have triggered great scientific interest to discover other 2D carbon allotropes. For example, following the success of graphene, penta-graphene,^7, 8^ phagraphene,^9, 10^ phographene,^11^ ψ-graphene,^12^ and θ-graphene^13^ have been intensively studied for exciting applications such as in nanoelectronics and energy storage. Most recently, a new 2D carbon allotrope, namely, popgraphene (standing for penta-octa-penta-graphene), has been proposed,^14^ which is composed of 5-8-5 carbon rings and has been theoretically demonstrated to possess a high dynamic, thermal, and mechanical stability. More importantly, thanks to its non-hexagonal ring structure, popgraphene exhibits extraordinary capabilities in the adsorption of lithium (Li) atoms, along with good conductivity and low energy barrier to Li diffusion, thus considered to be one of the most promising candidates for use in Li-ion batteries among 2D carbon allotropes.^12, 14-18^ Moreover, the unique structural and physical properties of popgraphene also promise potential applications in biomedical devices, such as highly sensitive biosensors and single-molecular DNA sequencers.^14, 19, 20^

Another advantage of popgraphene lies in the feasibility of its large-scale synthesis. For instance, in light of the controlled growth of atomically precise 5-8-5 carbon rings in graphene by simultaneous electron irradiation and Joule heating,^21^ it is natural to assume that large-area popgraphene that is comprised by ordered 5-8-5 carbon rings could also be synthesized. However, the above synthesis recipe employed an acceleration voltage of 80 keV for the incident electron beam,^21^ which falls in the range of the ejection threshold of carbon materials (i.e., 80-140 keV).^22^ Therefore, such process may increase the possibility of defect nucleation in the resultant popgraphene lattice, e.g., single vacancy (SV), double vacancy (DV), Stone-Wales defect (SW), as well as nanocracks resulted from defect aggregation or clustering, etc. Moreover, the 5-8-5 signature of the popgraphene lattice renders a diverse set of defect configurations, for example, there exist two configurations for SV and three configurations for DV. In light of the past research on 2D materials, defects would unavoidably disrupt the normal crystal lattice and play deterministic roles in the mechanical properties of 2D materials, which are critical metrics in the manufacturing, integration, and performance of devices on the basis of 2D materials.^23- 27^ Therefore, the existence of defects in popgraphene necessarily posts a concern of great relevance to the stability and durability of popgraphene-based devices.

In this regard, we assessed the existence of lattice defects and their effects on the fracture of popgraphene. Utilizing atomistic simulations, energetics of defect formation and the detailed fracture process of defective popgraphene have been examined. A comprehensive analysis of the fracture strength of popgraphene in the presence of defects was performed, and continuum models were employed to interpret the simulation results and to gain mechanistic insights. Key governing factors and mechanisms underlying the effect of defects have been identified. The present study provides a systematic set of data for understanding the fracture of defective popgraphene and offers critical guidance on the rational use of popgraphene in nanoscale devices.

## 2 COMPUTATIONAL METHOD

The structure of a popgraphene monolayer was illustrated in Fig. 1, being a 2D sheet comprising carbon atoms like graphene, but comprised by 5-8-5 carbon rings instead of the 6 rings in graphene. The unit cell of popgraphene was delimited in Fig. 1, which has 12 atoms enclosed by a conventional unit cell. The lattice constants were denoted as *c*_1_ and *c*_2_ . Similar to graphene, chirality can be defined for popgraphene. Here, chirality-dependent fracture behaviors were considered for the zigzag (ZZ) and armchair (AC) directions, where *σ_ZZ_* and σ_*AC*_ represent stress resulted from uniaxial tension normal to the ZZ and AC directions, respectively.

**Figure 1.**
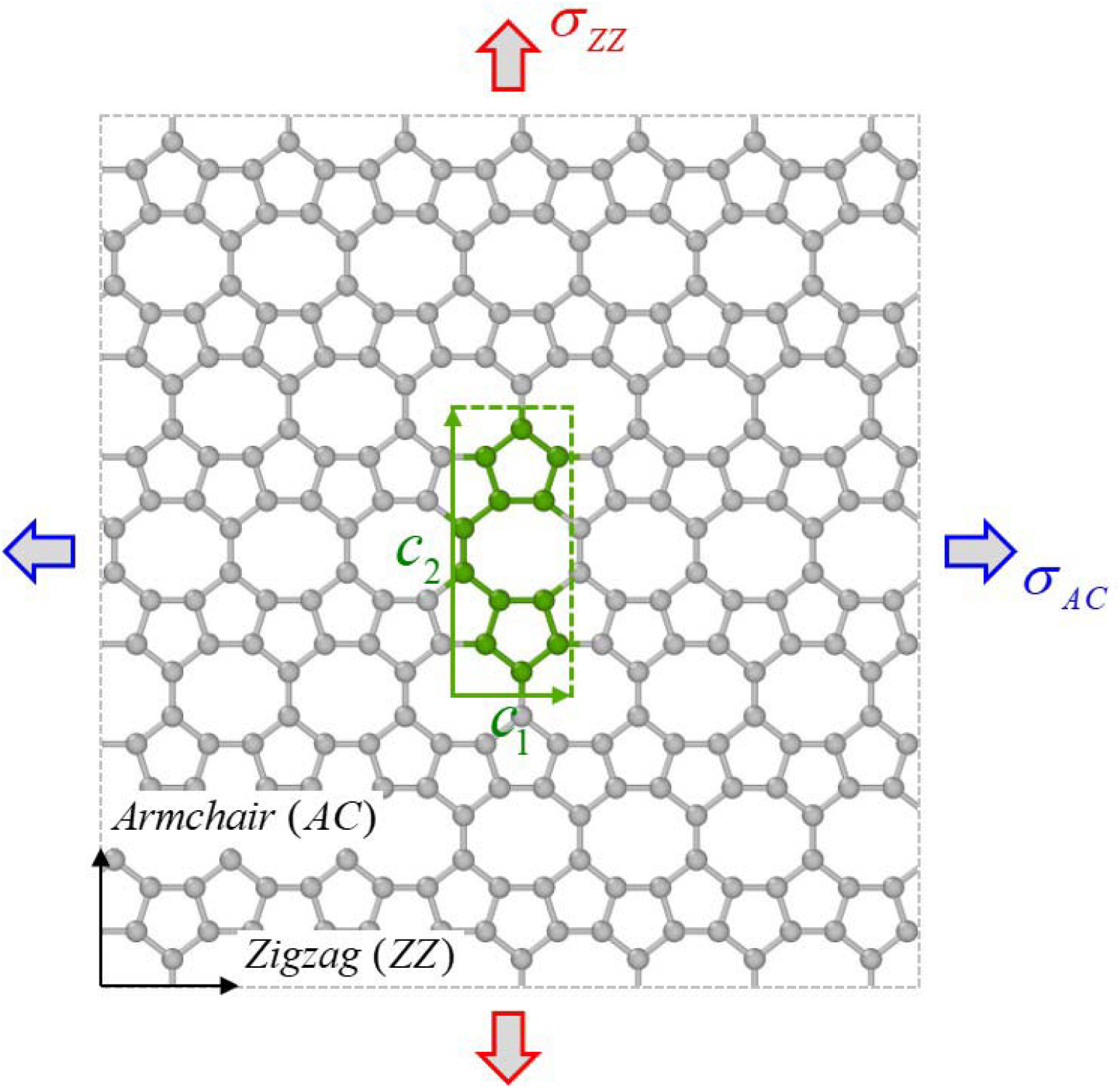
Supercell of the popgraphene monolayer with atoms enclosed by the olive dash rectangle representing the unit cell of the popgraphene structure. Stresses in popgraphene were denoted as *σ_ZZ_* and *σ_AC_* for uniaxial tension normal to the ZZ and AC directions, respectively.

Three scenarios of defect presence were considered in the popgraphene monolayer, including *i)* individual point defect (i.e., SV, DV, or SW defect); *ii)* distributed point defects (i.e., SV, DV, or SW defects with a range of defect densities); and *iii)* ZZ or AC nanocracks of different crack sizes. Below these three scenarios are elaborated in more details:

### i) Individual point defect

SV and DV were created by removing one and two atoms respectively from the popgraphene, while SW was formed by rotating a bond in popgraphene by 90°. The unique lattice sites where defects can form were illustrated in **Fig. 2a**. For SV, two defect configurations are possible, denoted as *SV_α_* and *SV_β_* as shown in **Fig. 2b**, while for DV and SW, each can have three defect configurations, denoted as *DV_αβ_*, *DV_βγ_*, and *DV_δη_*, and *SW_αβ_*, *SW_βγ_*, and *SW _βη_*, respectively, the atomic configurations of which were shown in **Fig. 2c-d**, respectively.

**Figure 2.**
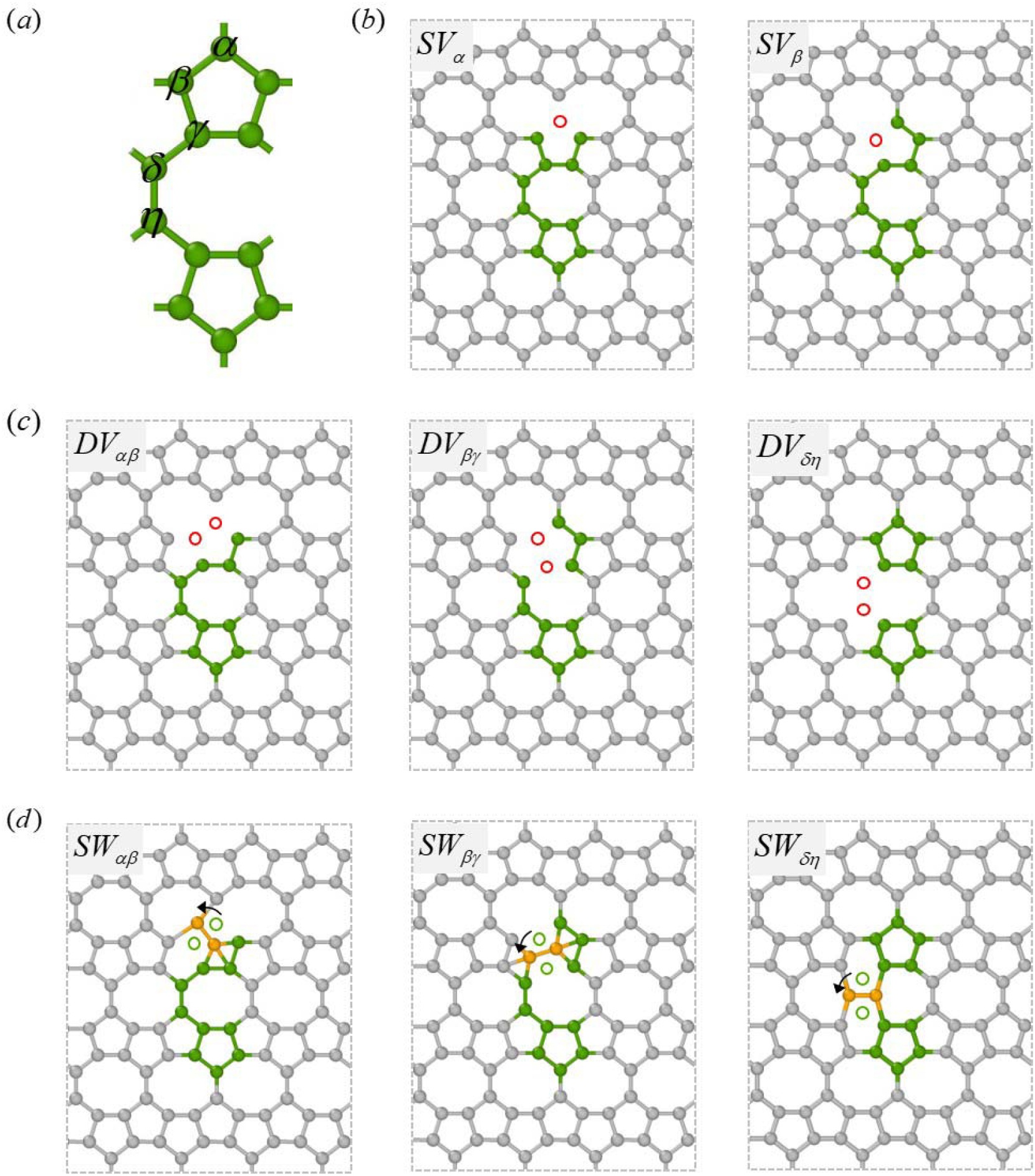
(a) Potential lattice sites where a point defect may form in popgraphene. (b) and (c) show the atomic configurations after the formation of SV and DV respectively, where the red open circles represent the atoms removed during the defect formation. (d) shows possible SW defect configurations with olive open circle and orange solid ball showing atoms before and after a 90° rotation required to create the SW defect, respectively.

### ii) Distributed point defects

Popgraphene with distributed point defects was created by assigning one of the three types of defects, i.e., SV, DV, or SW, of different densities into a popgraphene monolayer. The defect density ranges from 0.25% to 2.0% (with the percentage being the number of atoms removed for SV and half of the number of atoms removed/rotated for DV/SW over the total number of atoms in a pristine popgraphene) with a stepwise increment of 0.25% (i.e., in total eight defect densities considered). The defects were evenly distributed within the popgraphene sheet and each defect may randomly assume one of its possible configurations (see Fig. 2). In addition, defects were prevented from being in close vicinity to each other to avoid unrealistic large strain localization.^28^ **Figure 3a** presents a representative defective popgraphene containing distributed SW defects of 0.5% defect density, demonstrating the random spatial distribution and the defect assuming the three different configurations, i.e., *SW_αβ_* ,*SW_βγ_*, and *SW _βη_* . To ensure good statistics, for each defect density (of a particular point defect), four sets of models were created. The distributed defects cause out-of-plane crumping throughout the popgraphene to release the strain accumulated from the defects. **Figure 3b** showcased representative crumpled popgraphene configurations equilibrated at 1 K, showing an increased degree of crumpling with the increase of defect density.

**Figure 3.**
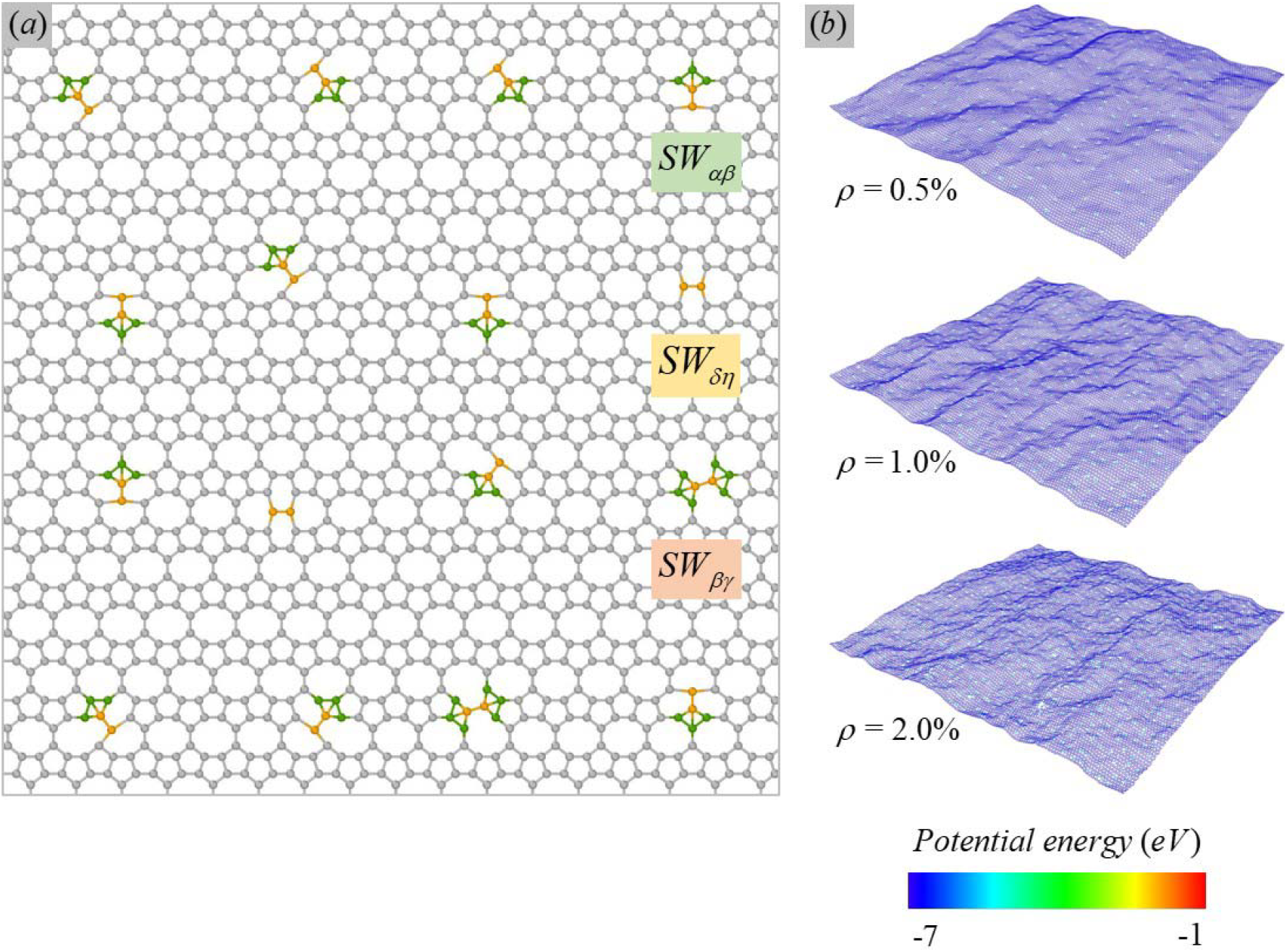
(a) A partial sheet within a representative defective popgraphene containing distributed SW defects of 0.5% defect density, where the defect may assume three different configurations of *SW_αβ_*, *SW_βγ_*, and *SW _βη_* before relaxation. (b) Defective popgraphene sheets containing distributed SW defects of defect density *ρ =* 0.5%, 1.0%, and 2.0%, after relaxation at 1 K, showing crumpling throughout the sheet.

### iii) Nanocracks

ZZ and AC nanocracks were considered in popgraphene as shown in **Fig.4a-b**, with the internal crack size denoted as 2*a* . The crack was formed by removing atoms from the pristine popgraphene sheet. Different crack sizes (i.e., *a* values) were considered in this study, ranging from 9.36 Å to 61.34 Å for ZZ cracks and 8.42 Å to 61.95 Å for AC cracks. Fracture energy and *J*-integral were computed for quantitative analysis of the nanoscale fracture of popgraphene. In particular, *J*-integral was calculated via the equivalent domain integral (EDI) method,^29^ the integration paths and domain of which were illustrated in **Fig. 4c**. The detailed calculation method of *J*-integral can be found below in **Section 3.4**.

**Figure 4.**
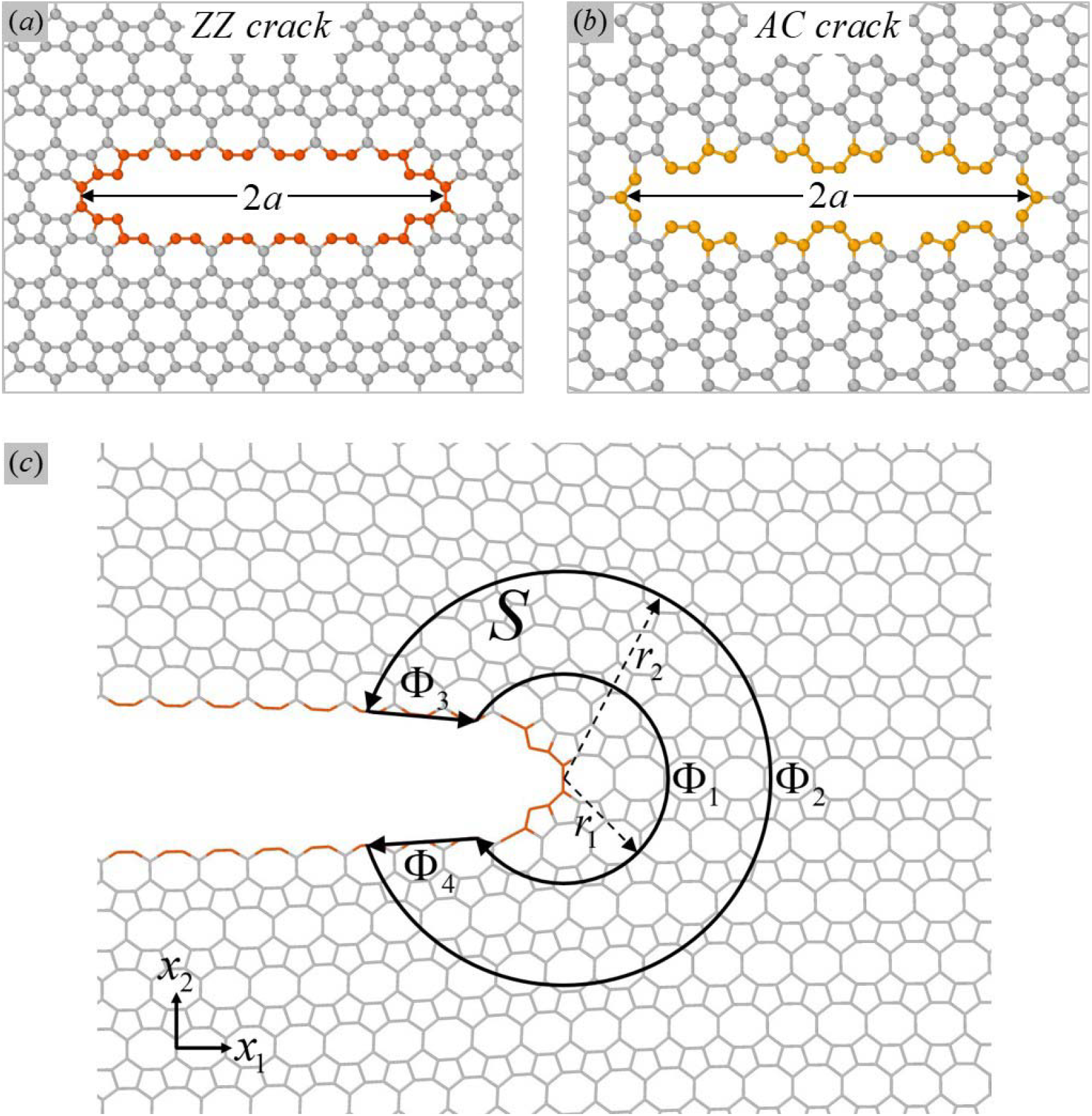
Atomistic configurations of (a) ZZ and (b) AC cracks in popgraphene monolayer with internal crack size denoted as 2*a* . (c) illustrates the integration paths and domain delineated around a crack tip for the calculation of *J*-integral, using a ZZ crack tip at the onset of crack growth as an example configuration. Colored atoms in (a-c) highlight atoms on crack surfaces.

Prior to studying the fracture behaviors of defective popgraphene, the energetics of those defects was examined by calculating their formation energy in popgraphene using first-principles calculations. The SIESTA package^30^ was employed, with the exchange-correlation functional being the generalized gradient approximation (GGA) parametrized according to Perdw-Burke- Ernzerh (PBE).^31^ Results were obtained using double-ζ polarized basis set, an energy cut-off of 250 Ry, and residual forces smaller than 0.03 eV/Å. The simulation supercell is fully periodic, has a dimension of 22.24 Å, 18.31 Å, and 15 Å for the ZZ, AC, and out-of-plane directions respectively, and contains 144 atoms for pristine popgraphene.

The subsequent fracture was studied through molecular dynamics (MD) simulations using the LAMMPS package,^32^ where the interaction among carbon atoms was described by the adaptive intermolecular reactive empirical bond order (AIREBO) potential.^33^ The equilibrium lattice constants of popgraphene, i.e., *c*_1_ and *c*_2_, calculated based on the potential were 3.71 Å and 8.90 Å respectively, being very close to 3.68 Å and 9.11 Å from first-principles calculations.^14^ It is worthy to note that the first nearest cut-off radius of the potential was modified from the original value of 1.7 Å to 2.0 Å to avoid the non-physical post-hardening behavior.^34- 36^ Periodic boundary conditions were kept for all directions of the simulation supercell. The popgraphene monolayer model used in the fracture study had an in-plane (i.e., ZZ×AC) dimension of 500.53 Å by 489.48 Å and 89,100 atoms, and the out-of-plane dimension was set at 500 Å to avoid interlayer interactions. After the creation of the defect, the popgraphene sheet was relaxed using the isothermal-isobaric (NPT) ensemble^37, 38^ at T=1 K for 400 picoseconds (ps) to ensure stress-free conditions in the in-plane directions prior to loading, and then deformed by a uniaxial tension of a strain rate of 1 ⨯ 10^-3^ ps^-1^ normal to the ZZ or AC direction till fracture. During the deformation process, the zero-pressure condition was maintained for the other in-plane direction normal to the loading direction. The timestep of 1 femtosecond was used for all simulations.

## 3 RESULTS AND DISCUSSION

### 3.1 Energetics of an individual point defect

Before studying the influence of defects on the fracture of popgraphene, it is important to first evaluate the formation energetics of those individual point defects in popgraphene. The formation energy, Ω, of an individual point defect can be defined as

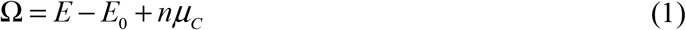

where *E* and *E*_0_ are the total energy of popgraphene with and without the defect, *n* is a prefactor, being 0 for SW defect, 1 for SV defect, and 2 for DV defect, and *μ_C_* is the chemical potential of a carbon atom in popgraphene, which was obtained from *E*_0_ divided by the number of atoms in the pristine popgraphene sheet. The formation energy data for defects in popgraphene together with those for defects in graphene are listed in **Tab. 1**. As noted from **Tab. 1**, the formation energies of the defect in popgraphene are generally comparable to those of graphene, and in some cases, e.g., *DV_βγ_* and *SW_δη_*, can be considerably smaller owing to the existence of more active lattice sites in non-hexagonal rings in popgraphene.^14^ Therefore, it is critical to study the fracture of defective popgraphene, the results of which will be presented in the following sections.

**Table 1.**
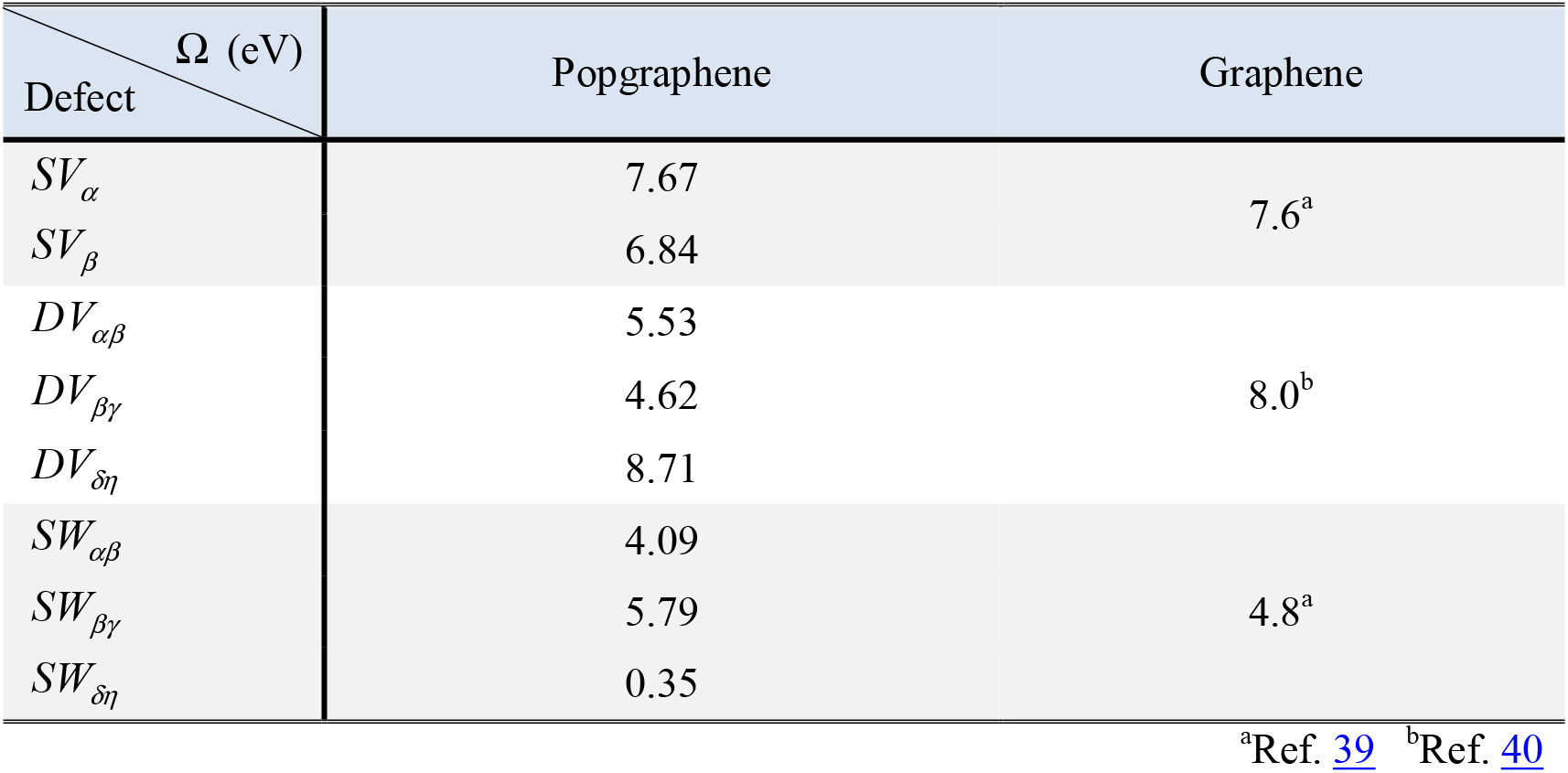
Formation energy of an individual point defect in popgraphene and graphene

### 3.2 Fracture of popgraphene with an individual point defect

The stress-strain response of pristine popgraphene with and without an individual point defect was plotted in **Fig. 5a-b** for uniaxial tensile strain normal to ZZ (i.e., σ_*ZZ-i*_, the subscript *i* indicates defect scenario *i*) and AC (i.e., *σ_AC-i_* ), respectively, which exhibits similar characteristics with that of graphene.^41^ Based on the above stress-strain curves, fracture stresses,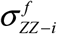 and 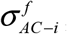, can be determined, as shown for each defect configuration in **Fig. 5c-d** and according to defect types in **Fig. 5e-f**. Overall, there is a relatively minor reduction in strength after an individual point defect was incorporated in popgraphene with the maximal reduction being 23% and 20% for 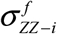 and 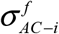, respectively, with strength reduction in the order of *DV* > *SV* > *SW* for both loading directions as shown in **Fig. 5e-f**. Meanwhile, comparing the strength of individual defect configuration within each defect group, for 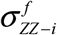, it reveals that *SV*_α_ < *SV_β_*, *DV_δη_* > *DV_βγ_* > *DV_αβ_*, and *SW_δη_* > *SW_αβ_* > *SW_βγ_*, while for 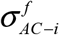, the trend is nearly opposite to that of 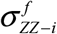 except that *DV_δη_* and *SW_δη_* keep being the largest among their respective defect type.

**Figure 5.**
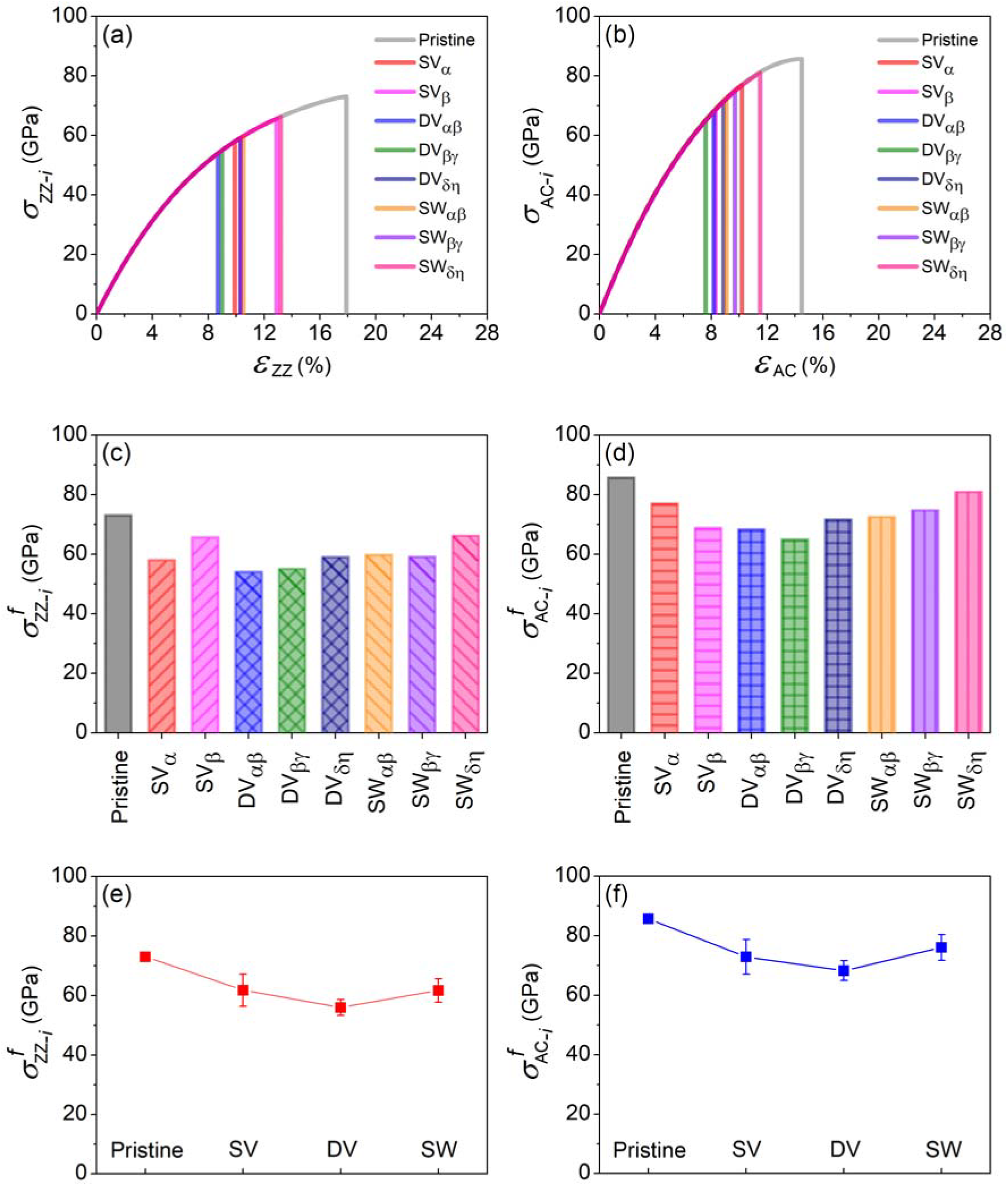
(a-b) Stress-strain curves and (c-d) fracture stresses of popgraphene with and without an individual point defect uniaxially loaded normal to the ZZ and AC directions, respectively. (e) and (f) plot the averaged fracture stresses calculated from (c) and (d) according to defect types, i.e., type of SV, DV, or SW. Error bars represent the standard deviation of the fracture stress calculated from each defect type.

To understand the factors contributing to the difference in fracture stress reduction, the atomic configurations and local stress states of individual defects at relaxed states prior to loading were plotted, as shown in Fig. 6. Examining those results, we identified two controlling factors responsible for the fracture initiation: *i) geometry of the defect*: larger defect or defect less symmetric with respect to the loading direction will experience higher stress concentration, causing lower fracture stress; and *ii) critical bond where fracture initiates*: the critical bond may exist within or away from the defect. Below for simplicity of discussion, we only focus on the loading normal to the ZZ direction (thus the 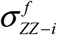 stress), as the loading normal to the AC direction (the 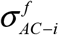 stress) follows the same line of reasoning. Specifically, for the SV group (see **Fig. 6a**), *SV_β_* exhibits a smaller effective flaw size than that of *SV_α_* and the critical bond for the case of *SV_β_* is farther from the defect than that for *SV_α_*, thus less stress concentration. This leads to higher fracture stress 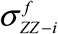 for *SV_β_* than that for *SV_α_*, as shown in **Fig. 5c**. For the DV group, as shown in **Fig. 6b**, *DV_δη_* has a more symmetric defect shape than that of *DV_αβ_* and does not have a dangling bond in the critical bond as *DV_βγ_* does, which consequently make *DV_δη_* the one with the largest 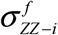 among the DV group. Comparing *DV_αβ_* with *DV_βγ_*, we see that despite having a dangling bond in its critical bond, *DV_βγ_* shows smaller defect size with the dangling bond slightly misaligned, which is a possible reason for a slightly larger 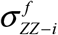 for *DV_βγ_* than that for *DV_αβ_* . For the *SW* group shown in **Fig. 6c**, we see that the fracture stress 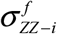 falls in the order of *SW_δη_* > *SW_αβ_* > *SW_βγ_*. This can be directly attributed to the stress concentration in the critical bond, which is in the order of *SW_δη_* < *SW_αβ_* < *SW_βγ_*.

**Figure 6.**
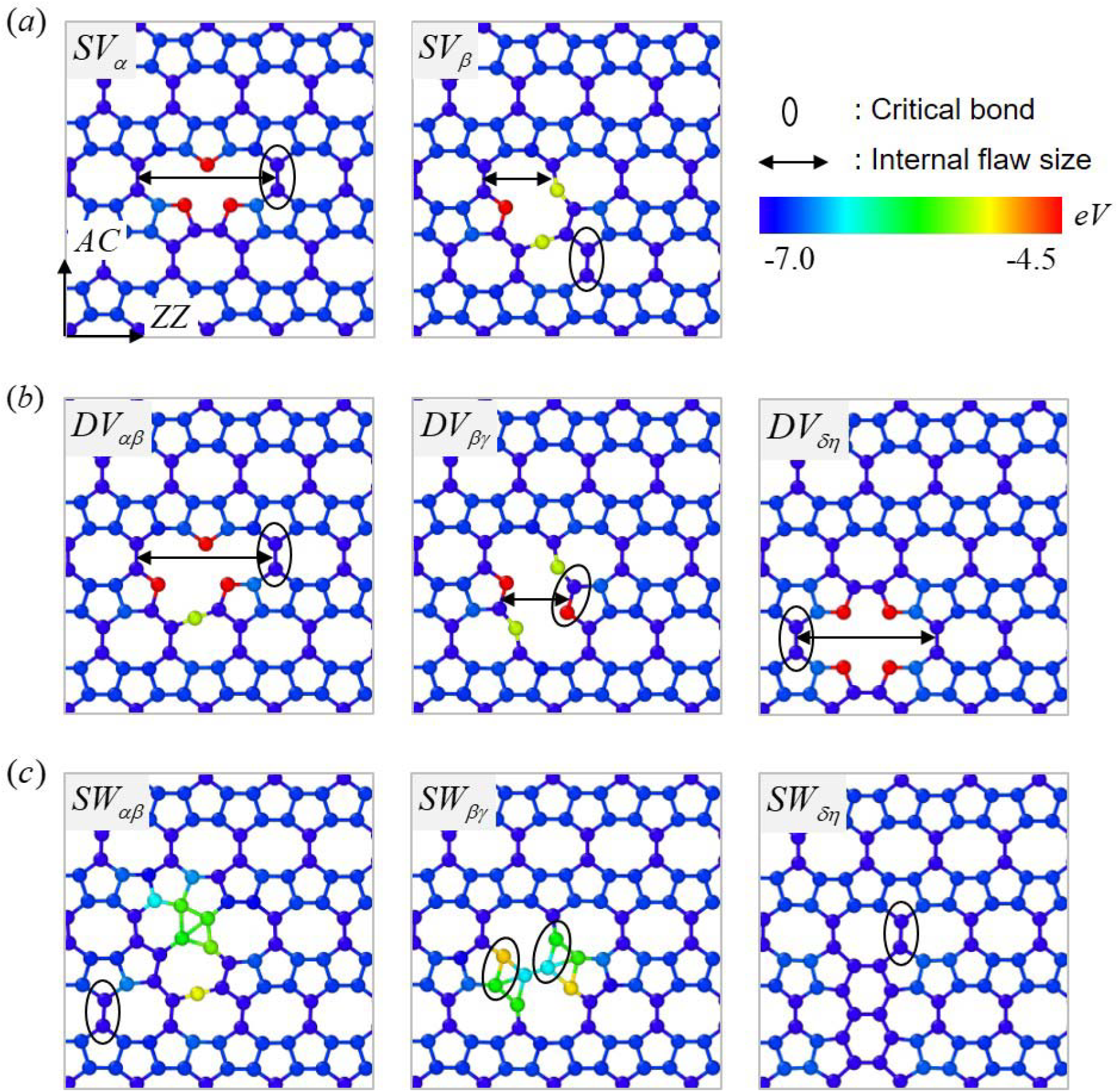
The relaxed atomic configuration of popgraphene prior to loading with individual (a) SV, (b) DV, and (c) SW defects. Atoms are colored according to potential energy contour with a scale bar from -7.0 to -4.5 eV. The critical bonds where the fracture occurs were enclosed by black ellipses, and the arrows indicated the effective flaw sizes associated with SV and DV defects under loading normal to the ZZ direction.

### 3.3 Fracture of popgraphene with distributed point defects

Fracture stresses,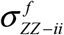 and 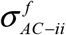 (the subscript *ii* indicates defect scenario *ii*), as functions of defect density for popgraphene monolayers with SV, DV, or SW defects were shown in **Fig. 7a-c** and **Fig. 7d-f**, respectively. It can be seen that for all three types of defects, the fracture stress along either direction is inversely proportional to the defect density, showing a nice linear trend. Note that a higher temperature than 1 K (temperature in our simulations), e.g., 300 K, would promote overall larger crumpling amplitude after the relaxation of the popgraphene monolayer. However, we have confirmed that the fracture stress of popgraphene loaded at 1 K is independent of the initial degree of crumpling [see **Electronic Supplementary Information (ESI) S1 for details**]. This finding is in agreement with Wei et al.’s study on polycrystalline graphene, where the initial out-of-plane deformation due to grain boundaries can be easily eliminated upon loading, thus having a very limited effect on the strength of the material.^26^ In addition, we observed that the reduction of strength is more pronounced with the existence of SW defects. It is also worth noting that the fracture stress for popgraphene with distributed SW defects shows considerable variation, particularly at large defect densities. This can be attributed to the high prestress localization in atoms around the SW transformation (see **ESI S2 for details**).

**Figure 7.**
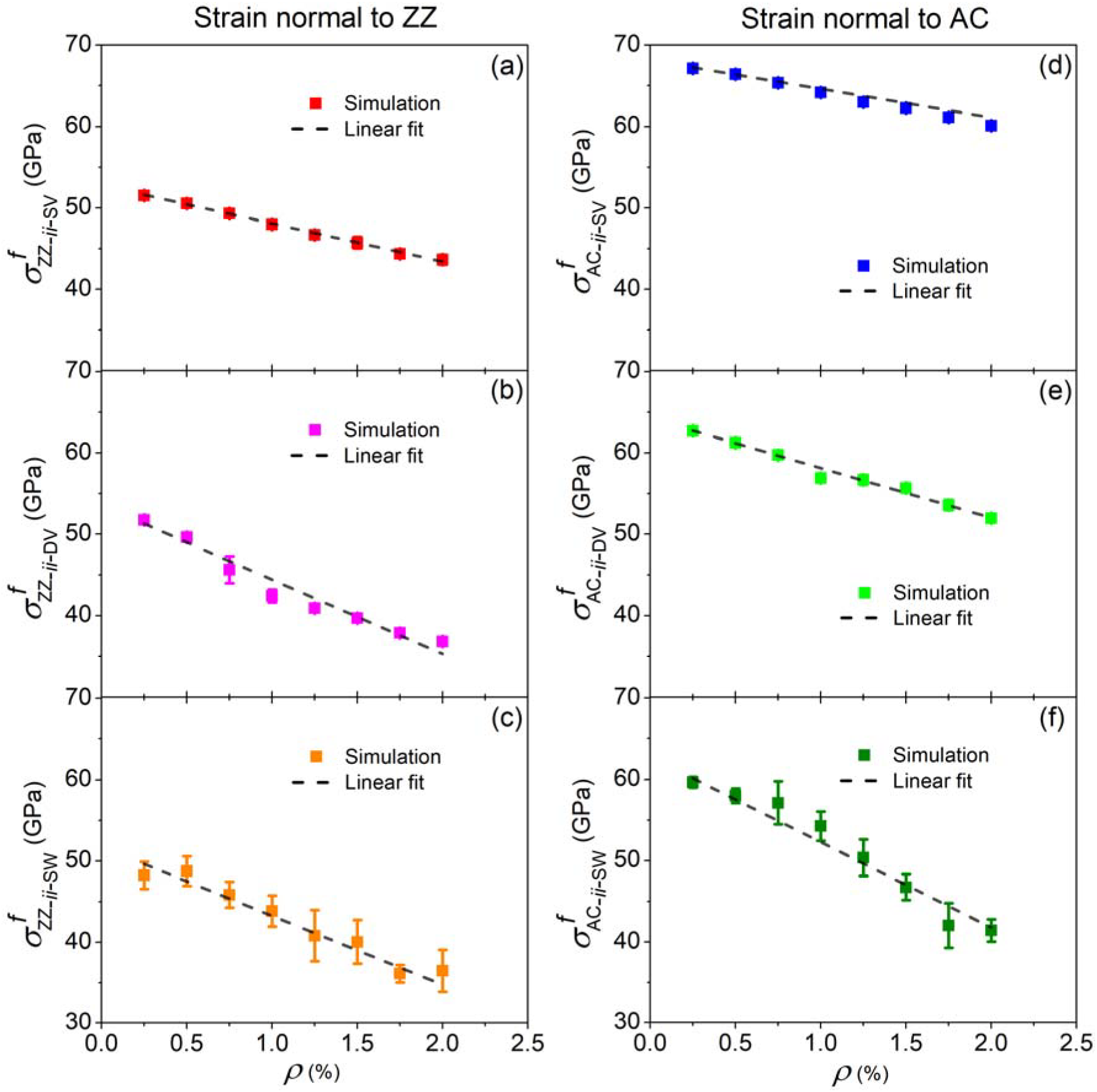
Fracture stresses, 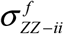 and 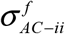, of defective popgraphene as functions of defect density *ρ* for distributed SV [(a) and (d)], DV [(b) and (e)], and SW [(c) and (f)] in popgraphene. Error bars are the standard deviation of the fracture stress calculated from the four sets of models.

### 3.4 Fracture of popgraphene with nanocracks

The MD simulated fracture stresses, 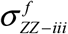 and 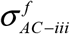 (the subscript *iii* indicates defect scenario *iii*), as functions of crack size *a* for popgraphene with ZZ and AC nanocracks were plotted in **Fig. 8a-b**, respectively. It can be seen that the fracture stress is decreasing with the increase of crack size, which is anticipated as a larger crack has a stronger stress concentration at the crack tip, and thus smaller fracture stress. Moreover, for cracks with the same crack size,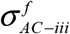 is found to be larger than 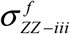. To further understand the fracture behaviors of popgraphene containing a nanocrack, we first examined the associated fracture energy,Γ_G_ ,^41^with the values for ZZ and AC cracks listed in **Tab. 2**. We see that Γ_*G*_ of AC cracks is larger than that of ZZ cracks, consistent with the order of their fracture stresses. Meanwhile, another important parameter that determines the nanoscale fracture behavior is lattice trapping, λ_*LP*_, defined as

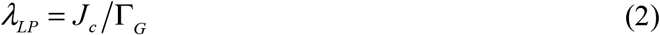
where *J_c_* is the critical *J*-integral calculated at the onset of crack, which can be calculated using the equivalent domain integral (EDI) method,^29^ with the discrete form of the domain integral (see domain *S* illustrated in **Fig. 4c**) given by

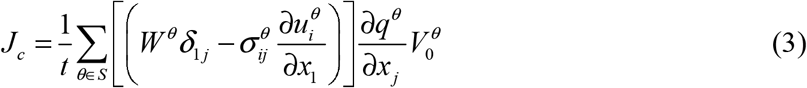

where *t* is the thickness of popgraphene sheet (assumed to be the same with that of graphene, i.e., 3.35 Å^1^), *W^θ^* is the strain energy density of atom *θ* calculated by the energy difference at the onset of crack growth and at relaxed state prior to loading, *δ*_1 *j*_ is Kronecker delta,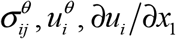 are stress calculated using the virial theorem,^41, 42^ displacement, and strain components (*i* = 1, 2 and *j* = 1, 2) at atom *θ*, respectively, 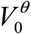 is the volume of atom *θ* before loading, and *q* is an arbitrary but continuous function given as

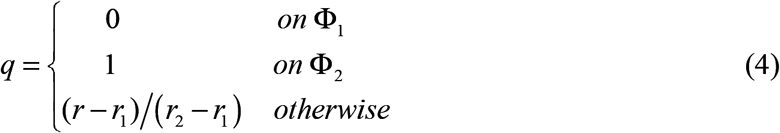

The strain 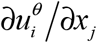was calculated according to the following equation^43^

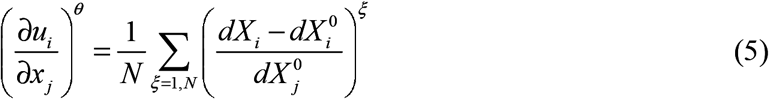
where *N* is the number of nearest neighbors of atom *θ* within the cut-off radius, i.e., 2 Å, of the potential function, *ξ* denotes the index of the neighboring atom,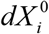, and 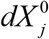 describes distances between atom *θ* and its neighboring atom *ξ*, specifically, distances after loading along the *i* direction, before loading along the *i* direction, and before loading along the *j* direction, respectively.

**Figure 8c-d** plotted the simulated lattice trapping, *λ_LP_*, as functions of crack size *a* for ZZ and AC cracks, respectively. We see that the simulated *λ_LP_* decreases with the increase of the crack size, consistent with the lattice trapping results from prior studies on graphene.^41, 44^ To enable a direct comparison of MD simulated fracture results with those from the continuum framework, the averaged lattice trapping was used,^44^ which are 1.04 and 1.06 for ZZ and AC cracks respectively as shown in **Fig. 8c-d**. Lattice trapping being close to unity indicates that the lattice trapping is negligible in popgraphene, thus allowing in directly using Griffith’s criterion to model the nanoscale fracture of popgraphene. Based on Griffith’s criterion, the critical fracture stress,σ_c_, is defined as^45^

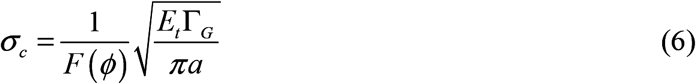
where *E_t_* is the tangent modulus of popgraphene calculated from the stress-strain curves of popgraphene at each corresponding crack size (see **ESI S3 for details**) to correctly reflect the nonlinear stress-strain response of popgraphene (as reflected in **Fig. 5a-b**),^44^Γ_G_and a were defined earlier in the text, and *F*(*ϕ*)is the geometry factor given by

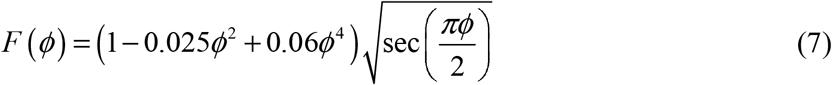
where *ϕ = W/*2*a* with *W* being the width of the popgraphene sheet (i.e., the width along *x*_1_ direction, see **Fig. 4c**). The predicted critical stress was plotted together with the simulated one in **Fig. 8a-b**. It is quite intriguing to see that the model prediction agrees very well with the MD simulated fracture stress for ZZ cracks, while underestimates that for AC cracks. To understand such discrepancy, we examined the detailed atomic process at the onset of fracture for ZZ and AC cracks as shown in **Fig. 9a-b**. We found that the ZZ crack shows a straight crack path while the AC crack undergoes a crack path deflection because the critical bond misaligns with the straight crack path (i.e., the bond in the ellipse in **Fig. 9b**). This crack deflection effectively renders the AC crack as loaded under mixed mode, and the fracture energy considering the deflection, Γ_D_, can be defined as ^46^

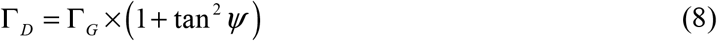

where *ψ* is the mixed mode angle, being 26.33° in the case of the AC crack based on **Fig. 9c**. The value of Γ_*D*_ was then obtained to be 12.05 *J/m*^2^, as listed in **Tab. 2**. Plugging in Γ_*G*_ in Eq. (6), We have the new prediction of the fracture stress for the AC crack

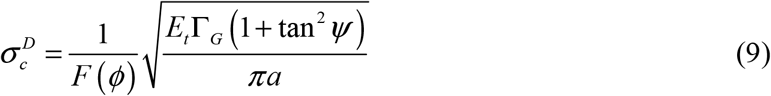

As seen in **Fig. 8b**, the new prediction from Eq. (9) shows excellent agreement with the simulation data. Note that Eq. (9) is reduced to Eq. (6) for the ZZ crack with the straight crack path. Furthermore, it is worth noting that Griffith criterion, in general, underestimates the critical fracture stress of very small cracks for both ZZ and AC cracks, e.g., at crack size *a* ≈ 10 Å in **Fig. 8a-b**. The reason is that such small crack may not be well regarded as a Griffith crack, which results in the inadequacy of the Griffith criterion.^25^

In addition, since the fracture of popgraphene was found to be brittle in nature, the fracture toughness is the true strength of popgraphene with engineering relevance. To this end, we calculated the fracture toughness of popgraphene. Based on the simulated critical stress 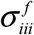 fracture toughness of popgraphene, *K_IC_*, can be extracted using Eq. (10) below

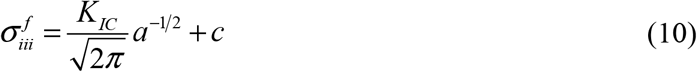

where *c* is a constant and its value is zero for a continuum solid. **Figure 8e-f** plotted 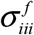 versus the inverse of the square root of the crack size *a* and the linear fit for the data for ZZ and AC cracks, respectively. From the slope of the linear fit, *K_IC_* can be calculated and its value is listed in **Tab. 2**. Comparing with graphene, the strength of popgraphene is only moderately weaker than graphene,^44, 47^ with a reduction of fracture toughness being 28% and 16% for ZZ and AC directions, respectively.

**Figure 8.**
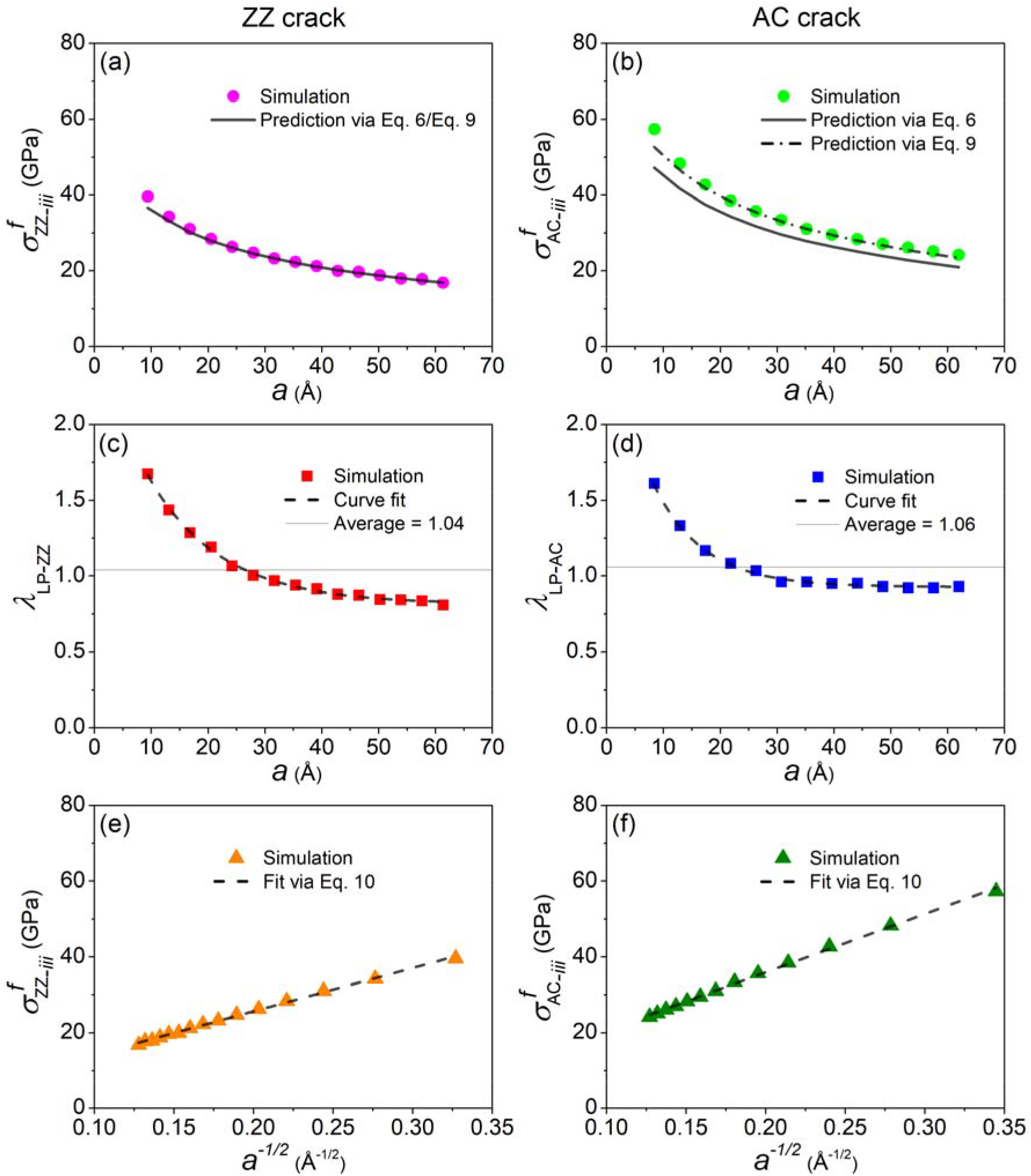
The simulated and predicted fracture stresses as functions of the crack size *a* of (a) ZZ and (b) AC cracks, respectively. Lattice trapping *λ_LP_* as functions of crack size *a* for (c) ZZ and (d) AC cracks, respectively. (e) and (f) plot the fracture stresses with respect to the inverse of the square root of crack size *a* .

**Table 2.** Fracture energy Γ and fracture toughness *K_IC_* of popgraphene and graphene

| Quantity \ Crack | Popgraphene |  | Graphene |  |
| --- | --- | --- | --- | --- |
|  | ZZ | AC | ZZ | AC |
| $\Gamma_G (J / m^2)$ | 7.54 | 9.68 | 10.49<sup>c</sup> (11.0<sup>d</sup>) | 11.27<sup>c</sup> (11.7<sup>d</sup>) |
| $\Gamma_D (J / m^2)$ | 7.54 | 12.05 | 11.0<sup>d</sup> | 15.9<sup>d</sup> |
| $K_{IC} (MPa\sqrt{m})$ | 2.87 | 3.86 | 4.0<sup>e</sup> | 4.57<sup>e</sup> |

<sup>c</sup>Ref. 27 <sup>d</sup>Ref. 45 <sup>e</sup>Ref. 44

**Figure 9.**
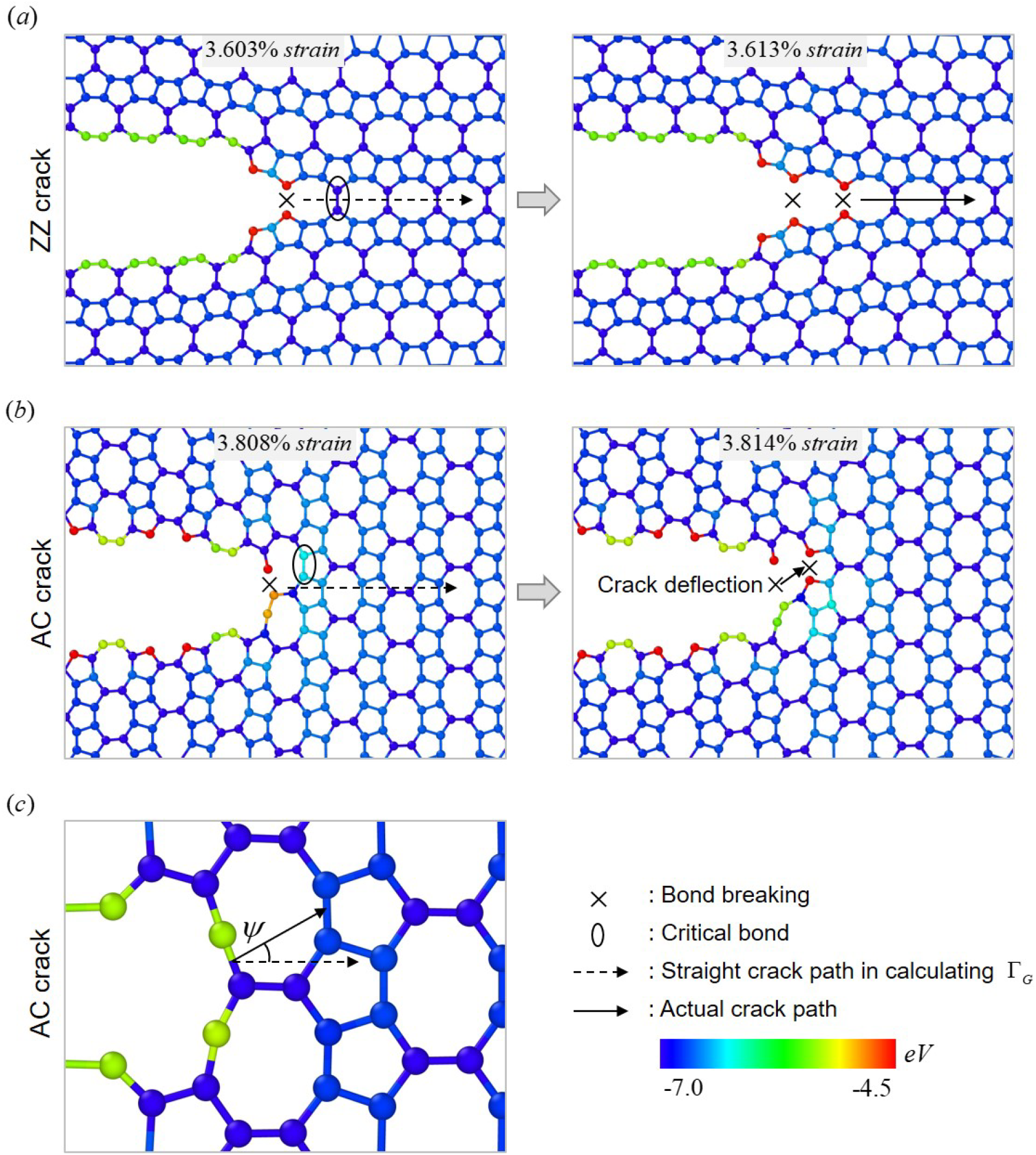
Atomic configurations colored by potential energy contour with a scale from -7.0 to - 4.5 eV at two consecutive strains immediately following the onset of fracture to show the critical bond breakage and the crack growth path for (a) a ZZ crack and (b) an AC crack. (c) illustrates the determination of the crack deflection angle *ψ* . Cross sign, ellipse, dash arrow line, and solid arrow line represent breaking of the critical bond, the critical bond, assumed crack path in calculating Γ_*G*_, and real crack path during a tensile test, respectively.

## 4 CONCLUSIONS

In summary, we systematically studied the fracture of defective popgraphene using MD simulations and continuum analysis. Three scenarios of defects were considered, including an individual point defect (SV, DV, or SW defect), distributed point defects of different defect densities, and nanocrack (AC or ZZ orientation) of different crack sizes. It was found that an individual point defect will not substantially deteriorate the strength of popgraphene and both the geometry of the defect (size and symmetry) and the critical bond where fracture initiates determine the fracture stress of popgraphene. When distributed point defects were present in popgraphene, the result showed that the fracture stress was inversely proportional to the defect density, showing a nice linear trend. Finally, for popgraphene containing a nanocrack, its fracture is brittle in nature and lattice trapping was found to be negligible for both ZZ and AC cracks. However, Griffith criterion can only accurately predict the fracture stress of popgraphene with ZZ cracks. It was further demonstrated that the disagreement between Griffith prediction and MD simulation for the AC crack was because Griffith criterion discarded the contribution of crack deflection in the AC crack. By introducing a mixed mode angle, Griffith criterion was modified to incorporate the crack deflection effect on the calculation of fracture energy to enable accurate prediction of the fracture stress of both ZZ and AC cracks. Overall, by comparing the tensile strength and fracture toughness of popgraphene with those of graphene, popgraphene possesses outstanding mechanical properties. Together with its unique structural and electronic properties, it is promising to explore popgraphene for novel nanoscale applications.

## ACKNOWLEDGMENTS

FM and JS acknowledge the financial support from Natural Sciences and Engineering Research Council of Canada (NSERC Discovery Grant # RGPIN-2017-05187) and from National Natural Science Foundation of China (NSFC Grant No. 51628101). The authors also thank Supercomputer Consortium Laval UQAM McGill and Eastern Quebec for providing computing power.

